# How Veeries vary: Whole genome sequencing resolves fine-scale genetic structure in a long-distance migratory bird, *Catharus fuscescens*

**DOI:** 10.1101/2023.07.25.550389

**Authors:** Abigail A. Kimmitt, Teresa M. Pegan, Andrew W. Jones, Kevin Winker, Benjamin M. Winger

## Abstract

Fine-scale resolution of spatial genetic structure is important for understanding a species’ evolutionary history and contemporary genetic diversity. For high-latitude species with high dispersal ability, such as long-distance migratory birds, populations typically exhibit little genetic structure due to high gene flow and recent postglacial expansion. Some migratory birds, however, show high breeding site fidelity, which might reduce gene flow such that population genetic structure could be detectable with sufficient genomic data. We sequenced over 120 low-coverage whole genomes from across the breeding range of a long-distance migratory bird, the Veery (*Catharus fuscescens*). As this species’ breeding range extends across both historically glaciated and unglaciated regions in North America, we evaluated whether contemporary patterns of structure and genetic diversity are consistent with historical population isolation in glacial refugia. We found strong evidence for isolation by distance across the breeding range, as well as significant population structure between southern Appalachian and northern populations. However, patterns of genetic diversity did not support southern Appalachia as a glacial refugium. Resolution of isolation by distance across the breeding range was sufficient to assign likely breeding origins of individuals sampled in this species’ poorly understood South American nonbreeding range, demonstrating the potential to assess migratory connectivity in this species using genomic data. Overall, our findings suggest that isolation by distance yields subtle associations between genetic structure and geography across the breeding range even in the absence of obvious historical vicariance or contemporary barriers to dispersal.

## Introduction

Resolving fine-scale genetic population structure in wild populations is important for understanding a species’ spatial and demographic evolutionary history as well as identifying microevolutionary processes underlying adaptation and population differentiation (Manel et al. 2003; Edwards et al. 2015; Lou et al. 2021). For species with high dispersal ability, however, resolving spatial genetic structure can be particularly challenging, as dispersal is associated with high levels of gene flow and minimal genetic structure (Slatkin 1987; Bohonak 1999; Claramunt et al. 2012; Medina et al. 2018). Seasonally migratory species, which often travel long distances between breeding and wintering grounds each year, are typically considered to have high dispersal, as their high vagility should reduce the impact of geographic barriers on their movement (Paradis et al. 1998; Medina et al. 2018; Everson et al. 2019; Claramunt 2021). Yet, in many bird species, long-distance seasonal migration is associated with limited dispersal between breeding sites, as adult migratory birds frequently exhibit high interannual fidelity to their breeding sites (Winger et al. 2019). By contrast, natal dispersal patterns remain poorly understood in small-bodied migratory birds. Breeding site fidelity and natal philopatry have the potential to reduce gene flow across the breeding range, such that long-distance migrants could still exhibit fine-scale genetic structure despite their long seasonal journeys and high dispersal potential. Here, we combine dense geographic sampling with whole-genome sequencing to investigate whether range-wide, fine-scale genetic structure can be resolved in the Veery (*Catharus fuscescens*), a Nearctic-Neotropical long-distance migratory songbird. We use these data to test patterns of genetic differentiation and diversity across the species’ range, evaluate the phylogeographic history of the species, and assess migratory connectivity between breeding and nonbreeding populations.

The Veery is an ideal species to test for the presence of fine-scale genetic population structure in migratory birds, given its long-distance migrations and high adult breeding site fidelity (Outlaw et al. 2003; Heckscher et al. 2011; Hobson and Kardynal 2015). The Veery breeds across wet forested habitats of the boreal and the temperate-boreal transition (‘hemiboreal’) belt, coastal forests of the northeastern US and Canada, the Appalachian Mountains, and riparian canyons in the mountains of western North America (Fig. 1; Heckscher et al. 2020). Previous work has identified five phenotypic subspecies based on subtle geographic population differences in plumage coloration (Phillips 1991; Pyle 1997). The only previous phylogeographic work on this species evaluated mitochondrial differentiation between the eastern and western extremes of the breeding range (Newfoundland versus Washington), identifying subtle but distinct genetic differentiation (Topp et al. 2013). Here, we employ dense, range-wide genomic sampling to investigate patterns of spatial genetic differentiation and evaluate the concordance of phenotypic subspecies descriptions with patterns of genetic differentiation. Given the potential for low genetic structure across the breeding range of this species, we tested for evidence of population genetic structure using low-coverage whole genome-sequencing (lcWGS). The development of cost-effective lcWGS allows inference based on orders of magnitude more loci than reduced representation genome sequencing (Lou et al. 2021), which might facilitate detection of subtle genetic patterns not otherwise evident (Novembre et al. 2008).

**Figure 1.**
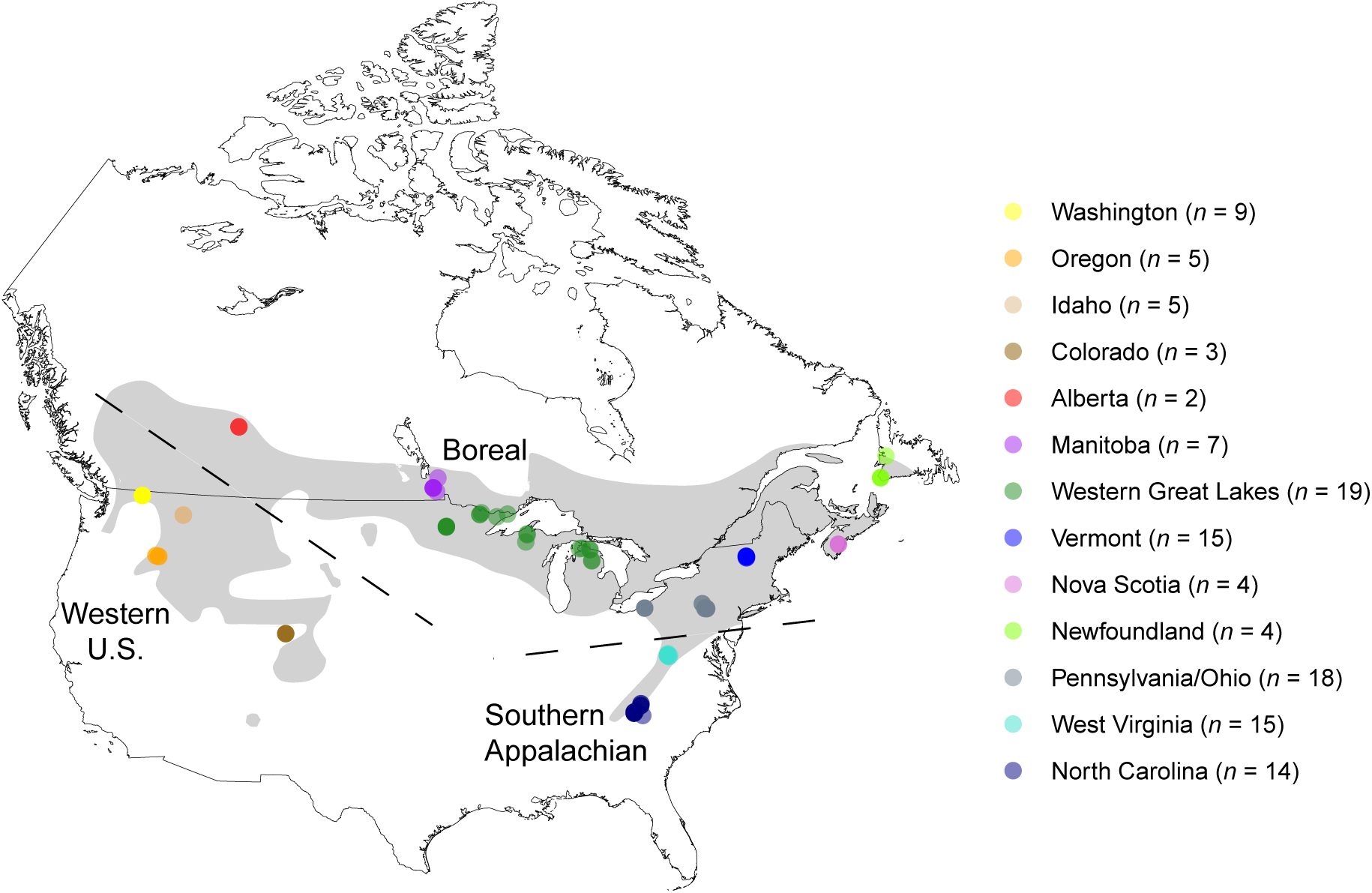
Map of sampling locations for the Veery (*Catharus fuscescens*). The approximate breeding range is highlighted in light gray (BirdLife International). Each point represents an individual, such that darker shading indicates multiple individuals. Dotted lines indicate approximate boundaries of the western U.S., boreal, and southern Appalachian populations identified by clustering analyses (Fig. 2). Four individuals sampled on their wintering grounds are not included in the map but were sampled in the Amazon Basin in Bolivia and Paraguay in October–November.

The nonbreeding distributions of this species, which is entirely within South America, is more poorly understood (Remsen Jr 2001; Heckscher et al. 2020). Veeries are known to exhibit intra-tropical movements during the overwintering period, as observed from geolocation data from populations breeding in Delaware (Heckscher et al. 2011) and British Columbia (Hobson and Kardynal 2015). Individuals spend the early portion of the northern winter in the Amazon basin south of the Amazon River in November–December, before moving northwest to a second wintering site likely in response to seasonal flooding patterns (Heckscher et al. 2011; Heckscher et al. 2015; Hobson and Kardynal 2015). With limited tracking data available, assessing migratory connectivity between breeding and nonbreeding ranges remains a challenge but is paramount to identifying the ecological and conservation links between stages in the annual cycle (Webster et al. 2002; Ambrosini et al. 2019). By producing the first detailed phylogeographic study of this species, a secondary goal of our study was to use samples from the winter range to identify nonbreeding birds’ breeding population of origin.

Contemporary breeding ranges of long-distance migratory birds, such as the Veery, are typically found at mid or high latitudes, such that Pleistocene glacial cycles presumably forced populations into fragmented habitat when ice sheets advanced (Hewitt 2004; Svenning et al. 2015). Resulting isolation of populations in putative glacial refugia is thought to have generated discrete population structure that is detectable in contemporary populations through measures of genetic diversity and heterozygosity (Bohonak 1999; Weir and Schluter 2004), for example as observed in several North American migratory bird species with molecularly distinct populations (e.g., Ruegg and Smith 2002; Barrowclough et al. 2004; Milá et al. 2007; Spellman and Klicka 2007; Manthey et al. 2011; van Els et al. 2012; Winker et al. 2023). Molecular signatures in multiple North American bird species have supported several glacial refugia— which might also have been occupied by the Veery—including south of the glaciers to the east (e.g., southern Appalachian Mountains) and west (e.g., southern Rocky Mountains), and offshore of Newfoundland (e.g., Grand Banks) (Hewitt 2004; Soltis et al. 2006). Therefore, we assessed contemporary genetic structure across the breeding range in light of historic processes associated with geographic isolation in different refugia versus population expansion from a single glacial refugium (Le Corre and Kremer 1998; Mimura and Aitken 2007; Meirmans 2012; Westram et al. 2013; Wahlsteen et al. 2023). If southern Appalachia and the western regions were historic glacial refugia for the Veery, we predict higher nucleotide diversity and heterozygosity in these populations given their likely long-term population stability as source populations for an expansion into post-glacial higher latitude habitat.

## Methods

### (a) Study system and sampling

We used 121 frozen or ethanol-preserved *C. fuscescens* tissue samples from our institutions’ museum collections or provided by other museum collections (Fig. 1; Table S1). We also included 3 blood samples from Newfoundland provided by the New York State Museum (Fig. 1; Table S1). Fieldwork by the authors was approved by our Institutional Animal Care and Use Committees and all relevant permitting authorities (see Acknowledgments). All samples were collected during the breeding season, except for 4 individuals that were collected on their wintering grounds in South America in October–November (Bolivia: n = 3, Paraguay: n = 1; hereafter, ‘nonbreeding birds’) that we included to assess migratory connectivity. Our sample size for nonbreeding birds is small but includes most nonbreeding tissue samples available in North American museum collections. Specifically, these samples represent four out of only six available tissue samples from the overwintering period that are published in a compendium of museum collections (www.vertnet.org). Given the sampling dates and locations, the four nonbreeding birds were likely collected on their first wintering site (Heckscher et al. 2011; Heckscher et al. 2015; Hobson and Kardynal 2015).

We extracted DNA using DNeasy Blood and Tissue Kits (Qiagen Sciences, Germantown, MD, USA). We prepared libraries for low-coverage whole genome sequencing using a modified Illumina Nextera protocol (Therkildsen and Palumbi 2017; Schweizer et al. 2021). We sequenced all libraries on NovaSeq (200 samples per lane) using services provided by the University of Michigan Advanced Genomics Core.

### (b) Data processing

We trimmed remaining adaptors and low-quality bases from demultiplexed data with AdapterRemoval v2.3.1 using the –trimns and –trimqualities options (Schubert et al. 2016). We also removed low-quality read ends using fastp v0.23.2 (Chen et al. 2018b) with the--cut_right option to mitigate the potential for batch effects arising from differences between sequencing runs (Lou and Therkildsen 2022). Following trimming steps, samples had a mean of 4.8x coverage of the genome (range= 2.59–28.38 billion bases; 2.3x–25.1x coverage).

We confirmed all samples were *C. fuscescens* by assembling and analyzing mitochondrial genomes as described in a previous study (Kimmitt et al. 2023). We used BLAST in Geneious (v. 2021.2.2) on at least one mitochondrial gene from each individual to confirm species identification. As a chromosome-assembled genome of *C. fuscescens* was not available, we aligned all samples to a reference genome of a close relative, *C. ustulatus* (GenBank assembly accession number GCA_009819885.2bCatUst1.pri.v2, coverage = 60.58x) using bwa mem (Li and Durbin 2010) and Samtools (Li et al. 2009). We removed overlapping reads using clipOverlap in bamUtil (Jun et al. 2015), marked duplicate reads with MarkDuplicates, and assigned all reads to a new read group with AddOrReplaceReadGroups using picard (http://broadinstitute.github.io/picard/). All bam files were then indexed using Samtools (Li et al. 2009). The mean mapping rate across all samples used in analyses was 97.43% (range 93.98– 98.43%). We then used GATK v3.7 (Van der Auwera et al. 2013) to re-align samples around indels by applying RealignerTargetCreator to the entire dataset and using IndelRealigner for each sample.

We used ANGSD v0.9.40 (Korneliussen et al. 2014) to calculate genotype likelihoods using the GATK model from low-coverage sequencing data. Given the genotype uncertainty associated with low-coverage sequencing, all results were analyzed in a genotype likelihood framework, as this method uses probability-based inference to account for sequencing error (Korneliussen et al. 2014; Lou et al. 2021). Parameters used for each ANGSD analysis are described further below or detailed in Table S2.

### (c) Population structure

We calculated genotype likelihoods for all sites with a SNP *p*-value < 0.05 across the entire genome using ANGSD (Table S1). We then filtered mis-mapped or paralogous SNPs out of the dataset using ngsParalog v1.3.2 (https://github.com/tplinderoth/ngsParalog; Linderoth 2018). ngsParalog is designed for low-coverage sequencing data and implements a likelihood method to find mapping problems.

We used PCAngsd v1.10 (Meisner and Albrechtsen 2018) to conduct Principal Component Analyses (PCA) to visually assess spatial genetic structure. As PCA can be sensitive to genomic inversions that could obscure geographic structure (Novembre et al. 2008; Tian et al. 2008; Novembre and Peter 2016), we first ran PCAngsd separately for each chromosome using all 124 samples to identify possible inversions. We observed at least six chromosomes with evidence of clustering associated with putative inversions, so we analyzed each chromosome further for inversions using lostruct (Li and Ralph 2019) as implemented using PCAngsd with scripts available from https://github.com/alxsimon/local_pcangsd. We removed all microchromosomes with evidence of inversions (*n* = 8). We also removed all sex chromosomes from our dataset. For the remaining chromosomes, we then ran PCAngsd with the --admix option to estimate admixture proportions using a non-negative matrix factorization algorithm so that we could produce genome-wide PCAs and admixture plots. Two individuals sampled from Nova Scotia had an aberrantly high PCA covariance (> 0.2) such that they were visual outliers on the PCA (see Fig. S1); therefore, we excluded one of these individuals from the final PCAs to better facilitate visual assessment of range-wide structure patterns.

We implemented the *find.clusters* function from the R package *adegenet* (Jombart et al. 2010) using the covariance matrix produced by PCAngsd; *find.clusters* runs successive *K*-means with an increasing number of clusters (*K*) and then performs a goodness of fit analysis (BIC) to identify the optimal *K*. We also used a Mantel test in the *ade4* package v. 1.7–19 (Thioulouse and Dray 2007), with 1000 permutations, to determine whether genetic distance (using the proxy 1 – PCA covariance; Novembre et al. 2008) varied significantly with geographic distance between samples.

We calculated pairwise *F_ST_* between distinct populations. Specifically, we conducted analyses on three distinct populations that were revealed by the PCA-based clustering methods mentioned above (see also Results and Figs. 1-2): (1) “western” (i.e., western United States including Washington, Oregon, Idaho, and Colorado; *n* = 22), (2) “southern Appalachian” (i.e., West Virginia and North Carolina, *n* = 29), and (3) “boreal” (i.e., Canada from Alberta to Newfoundland, Western Great Lake states and Northeastern United States; *n* = 69) (Fig. 1). To make our *F_ST_* calculations computationally tractable, we randomly downsampled the reference genome, including SNPs and invariant sites, to create a set of loci that consisted of stretches of 2 kb loci at least 10 kb apart (yielding approximately 12% of the whole genome). We used scripts modified from https://github.com/markravinet/genome_sampler and excluded loci from regions removed by the inversion filters. We removed sites flagged by ngsParalog from the subsampled dataset and stored the loci in a BED file. We generated a site allele frequency (SAF) file in ANGSD with the -doSaf parameter and -sites filter to include only subsampled loci. We used winsfs (Rasmussen et al. 2022) to create 2-dimensional (2D) site frequency spectra (SFS) between each population pair. We then used the *F_ST_* index and stats function with the option - whichFst 1 (i.e., Bhatia estimator) in ANGSD to calculate pairwise *F_ST_* between unbalanced sample sizes.

**Figure 2.**
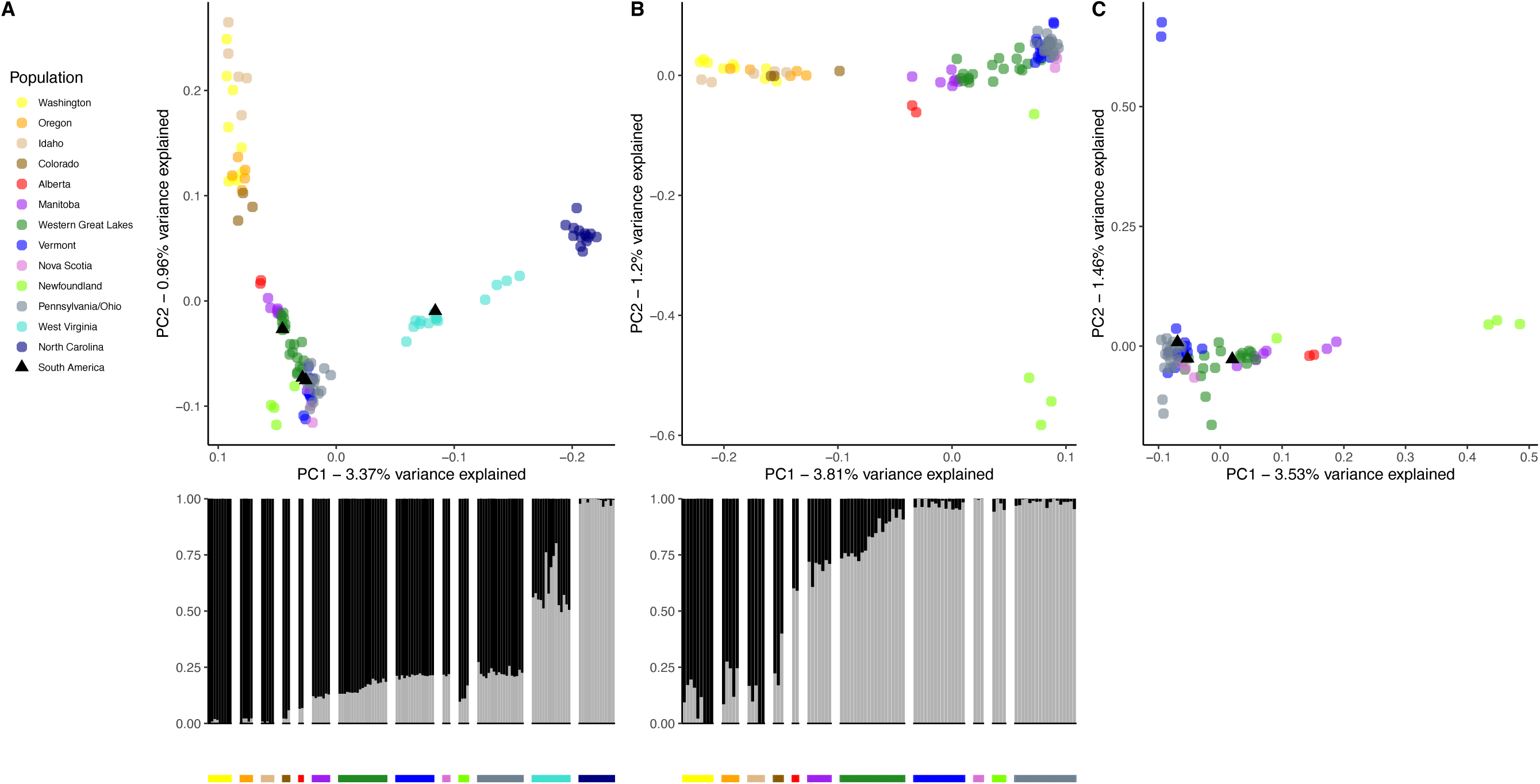
Principal Component Analysis (PCA) and admixture plots (*K* = 2) for (A) all samples and (B) all samples excluding the southern Appalachian samples (i.e., excluding West Virginia and North Carolina). (C) PCA of boreal and northern Appalachian samples only. Nonbreeding birds collected in the South America are displayed on Panel A and Panel C as black triangles to identify potential population of origin. The x-axis in panel A is reversed so that points are displayed west to east. The U-shaped curve shown in panel A, the nested genetic structure in Panel B and C, and the level of admixture across the geographic range suggests isolation by distance across the species’ range. The southern Appalachian (West Virginia and North Carolina) were supported as a distinct population from all other samples (A), and the western samples (Washington, Oregon, Idaho and Colorado) were supported as a distinct population when southern Appalachian samples were removed (B). By contrast, all boreal samples were identified as a single population but with evidence of isolation by distance.

Finally, to assess the direction of gene flow between the three populations, we calculated a directionality index (*ψ*) from the 2D SFS with a custom script from (Adams et al. 2023) using equation 1b from (Peter and Slatkin 2013). Balanced sample sizes are necessary to calculate *ψ* (Peter and Slatkin 2013), such that we randomly selected 22 individuals from both the southern Appalachian and the boreal populations to created new SAF and 2D SFS files between each population pair.

### (d) Genetic diversity and heterozygosity

For each of the three populations identified by the clustering analysis above (western, boreal, and southern Appalachian), we estimated population-level summary statistics for genetic diversity from the subsampled loci using ANGSD and winsfs. Since the sample size of the more geographically expansive boreal population was much larger than the other two populations, we randomly selected 30 individuals from the boreal population for population-level genetic diversity analyses. We generated a SAF using only subsampled loci in ANGSD that excluded flagged sites by ngsParalog and microchromosomes with detected putative inversions. We used winsfs to produce and folded a population-level 1-dimensional (1D) SFS. We then calculated pairwise *θ_π_* for each chromosome separately using the saf2theta and thetaStat functions in ANGSD. We compared *θ_π_* by chromosome among populations using a one-way analysis of variance (ANOVA). For each population we also calculated the total pairwise π / total number of sites across all chromosomes.

We also estimated individual-level heterozygosity. We created individual-level SAF files with the same subsampled loci used in genetic diversity estimates. We then used these SAF files with winsfs to generate individual 1D SFS. We calculated individual heterozygosity as the number of polymorphic sites divided by the total sites in each individual’s 1D SFS (Kersten et al. 2021). Exploratory analyses suggested that samples with very low (<4x) genomic coverage exhibited low individual-level heterozygosity relative to samples above 4x coverage. Therefore, we filtered all samples with less than 4x coverage (*n* = 35) out of the dataset for this analysis (retaining samples sizes of *n =* 15 for western U.S., *n =* 56 for boreal, and *n =* 14 for Southern Appalachian regions). We then compared population differences in individual heterozygosity using ANOVA and Tukey multiple pairwise comparisons. Since the three populations span large geographic ranges, we also tested for the presence of gradients of heterozygosity across latitude or longitude using linear models: 1) a western gradient across latitude including samples from Alberta and the western population, 2) a boreal gradient across longitude, and 3) an eastern gradient across latitude including samples from the southern Appalachian population as well as PA, OH, VT, Nova Scotia, and Newfoundland (Fig. 1).

## Results

### Population structure

The PCA pattern indicated isolation by distance, as the shape of the PCA reflects a sinusoidal curve typical of continuous structure (Fig. 2A; Novembre and Stephens 2008). The Mantel test confirmed isolation by distance, as genetic distance (1 – PCA covariance) was positively associated with geographic distance (Pearson’s correlation coefficient, *r* = 0.38, *p* = 0.001) using samples across the full breeding range (Fig. 3A). We also visually inferred two distinct population clusters in the range-wide dataset of breeding individuals (*n* = 120) in the PCA analysis, such that individuals from the southern Appalachian sampling regions (North Carolina and West Virginia) clustered separately from all other individuals, demonstrating a genetic break between the southern Appalachian samples and the northern Appalachian samples (i.e., Pennsylvania, Ohio, and Vermont; Fig. 2A). We confirmed that two clusters were the best fit for the data using the *find.clusters* tool. The admixture plot (*K* = 2) showed a gradual shift in population ancestry across the geographic range.

**Figure 3.**
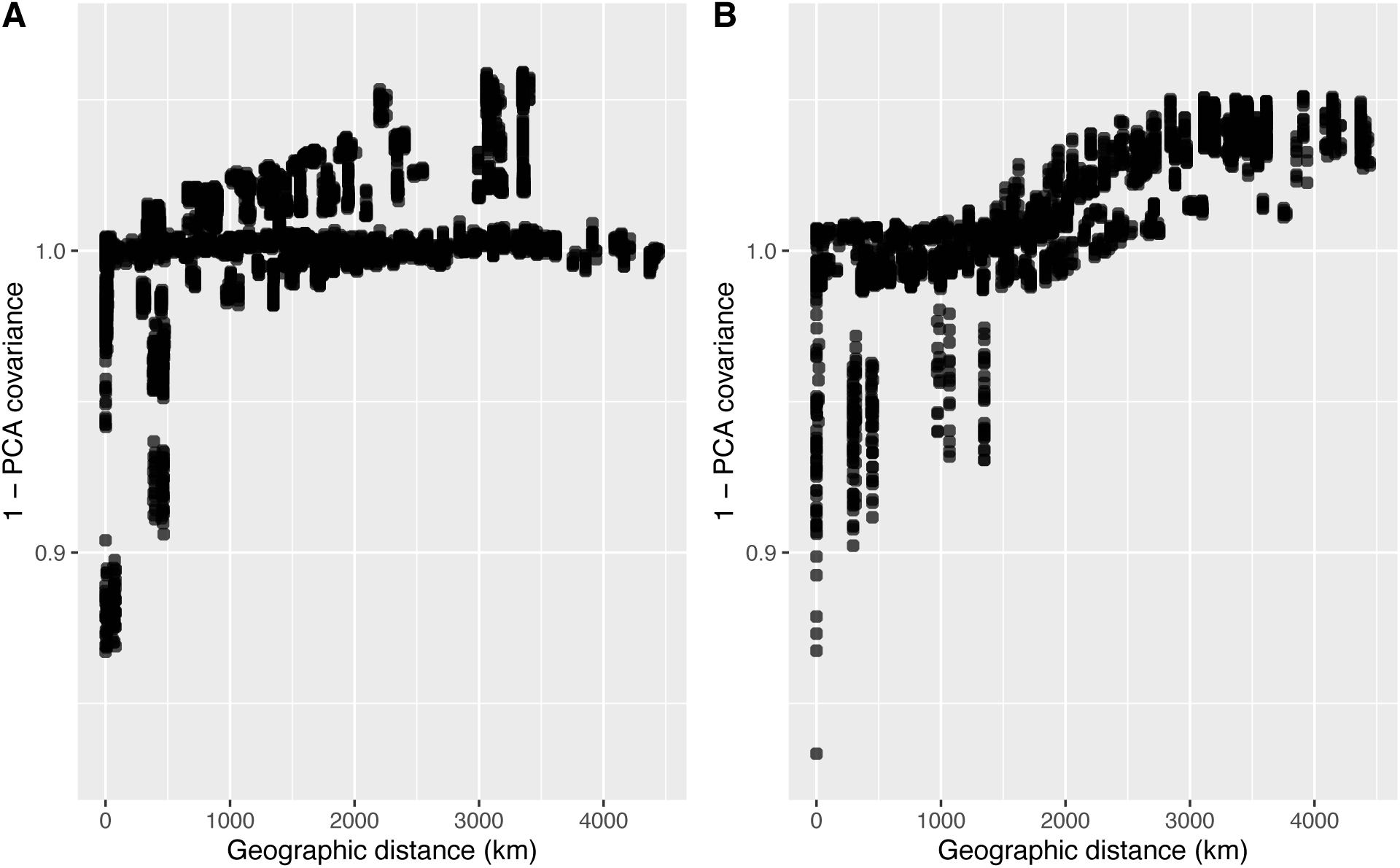
Genetic distance (as measured by 1– PCA covariance) between pairs of individuals is significantly positively correlated with geographic distance (A) using the range-wide dataset (corresponding to panel A in Fig. 2) and (B) the boreal, northern Appalachian, and western US sampling location data (corresponding to panel B in Fig. 2). The correlation recovered from the Mantel test is stronger in the subset of data in panel B (Pearson’s correlation coefficient, *r* = 0.78) compared to the range-wide data (Pearson’s correlation coefficient, *r* = 0.38) because of distinct population structure associated with the southern Appalachian population.

We next removed Appalachian samples from the dataset to determine if we could detect finer-scale genetic structure in the more genetically similar samples from western and boreal sampling regions. In the boreal and western populations dataset, the relationship between genetic distance and geographic distance was stronger than in the range-wide dataset (*r* = 0.78, *p* = 0.001; Fig. 3B). Without the southern Appalachian samples included, we also found that the western population was distinct from a boreal population, which also includes the northern Appalachian samples (Fig. 2B) and confirmed that two clusters was the best fit for this subset of the data using *find.clusters*. Finally, we ran a PCA on the boreal population samples alone to determine if subpopulations would be detectable on a further reduced geographic scale. We visually noted that 3 out of 4 of the samples from Newfoundland sorted separately on the PCA, suggesting that this isolated population could be distinct from other boreal populations. However, *find.clusters* did not assign distinct clusters associated with geography within the boreal population samples, consistent with the observed overlap among sampling regions within the PCA (Fig. 2). Finally, the relationship between genetic distance and geographic distance was weakest in the boreal population samples only (*r* = 0.22, *p* = 0.001).

Based on the PCA and clustering results, we conducted analyses of population differentiation and genetic diversity (next section) using three identified populations across the sampling regions (Fig. 1): southern Appalachian (i.e., WV, NC), western US (i.e., WA, OR, ID, and CO; hereafter the ‘western’ population), and the boreal belt (Alberta to Newfoundland) including the northern Appalachians (i.e., PA, OH, VT), hereafter the ‘boreal’ population. Pairwise *F_ST_* values were < 0.02 between all three populations, indicating low levels of population differentiation. Weighted pairwise population-level *F_ST_* was 0.008 between the boreal and southern Appalachian population and 0.006 between the boreal and western populations. *F_ST_* was highest between the southern Appalachian and western populations (0.014).

Finally, the directionality index was low (*ψ* < 0.03) for all pairwise comparisons: *ψ* = 0.002 (*z =* 0.01) between the southern Appalachian and western populations, *ψ* = -0.03 (*z =* - 1.04) between the boreal and western population, and *ψ* = 0.02 (*z =* 0.95) between the southern Appalachian and boreal populations. The direction of *ψ* might indicate that the boreal population has been a source population for expansion; however, these values were not significantly different from zero, supporting an isolation by distance model or a population expansion model in which populations are equidistant from the origin of expansion and are exhibiting comparable levels of gene flow between populations (Peter and Slatkin 2013; Adams et al. 2023).

### (b) Genetic diversity and heterozygosity

Neither nucleotide diversity (pairwise *θ_π_*) estimated per chromosome nor individual-level heterozygosity differed significantly between populations (Table 1; Fig. 4A: *F_2,87_* = 0.02, *p* = 0.982; Fig. 4B: *F_2,81_* = 1.09, *p* = 0.341). Across a latitudinal gradient in the montane west, heterozygosity scaled positively with latitude (Fig. 5A). Across a longitudinal gradient in the boreal belt, western samples had significantly higher heterozygosity than eastern samples (Fig. 5B). Finally, there were no significant latitudinal differences across the eastern latitudinal gradient from North Carolina to Newfoundland (Table 2; Fig. 5C).

**Figure 4.**
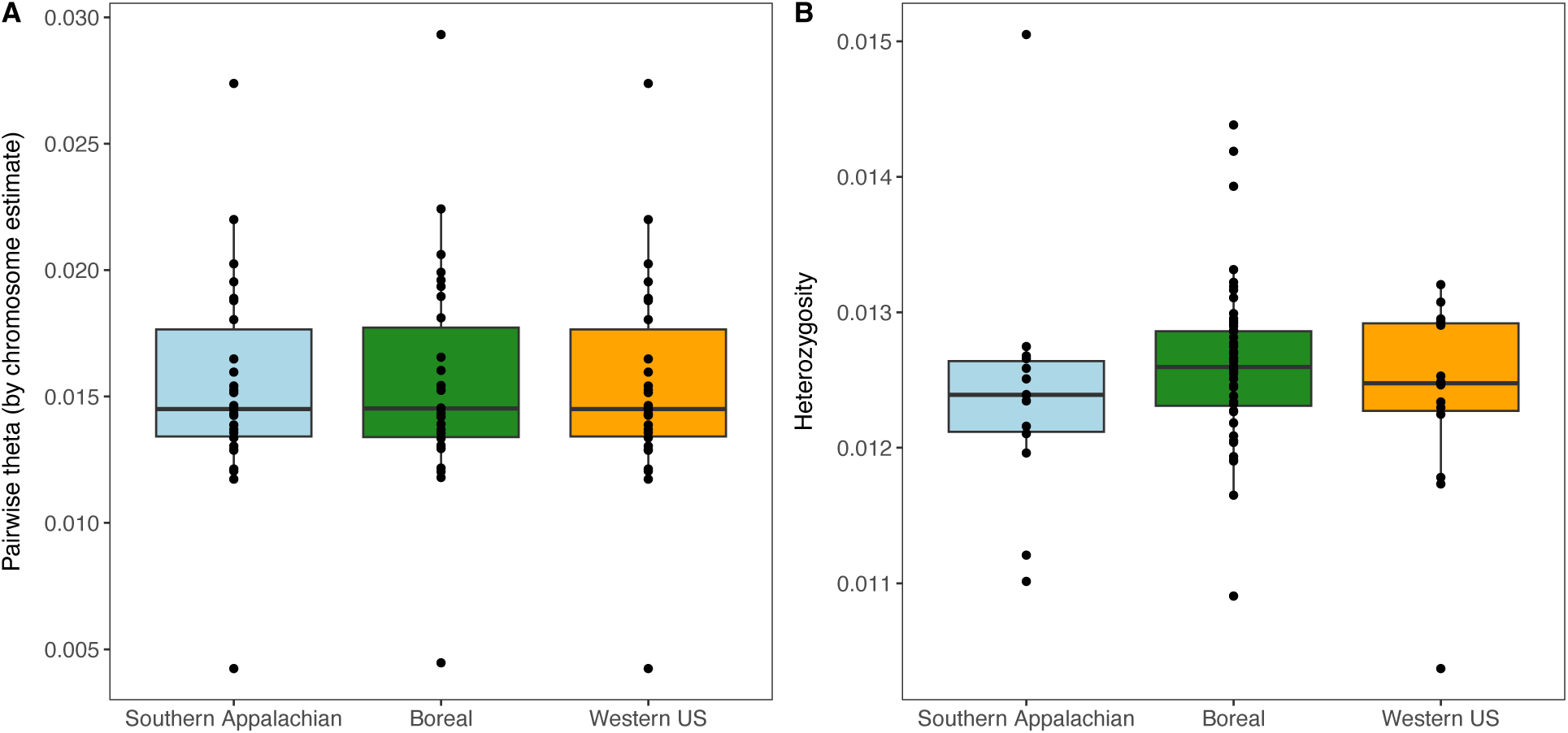
Comparisons of (A) genetic diversity as measured by pairwise *θ_π_*, and (B) individual heterozygosity of three populations of *Catharus fuscescens.* Population-level pairwise *θ_π_* was calculated per chromosome from population-level 1D site frequency spectra. Individual heterozygosity was calculated as the number of polymorphic sites by the total sites in the individual-level 1D site frequency spectrum. See also Table 1.

**Figure 5.**
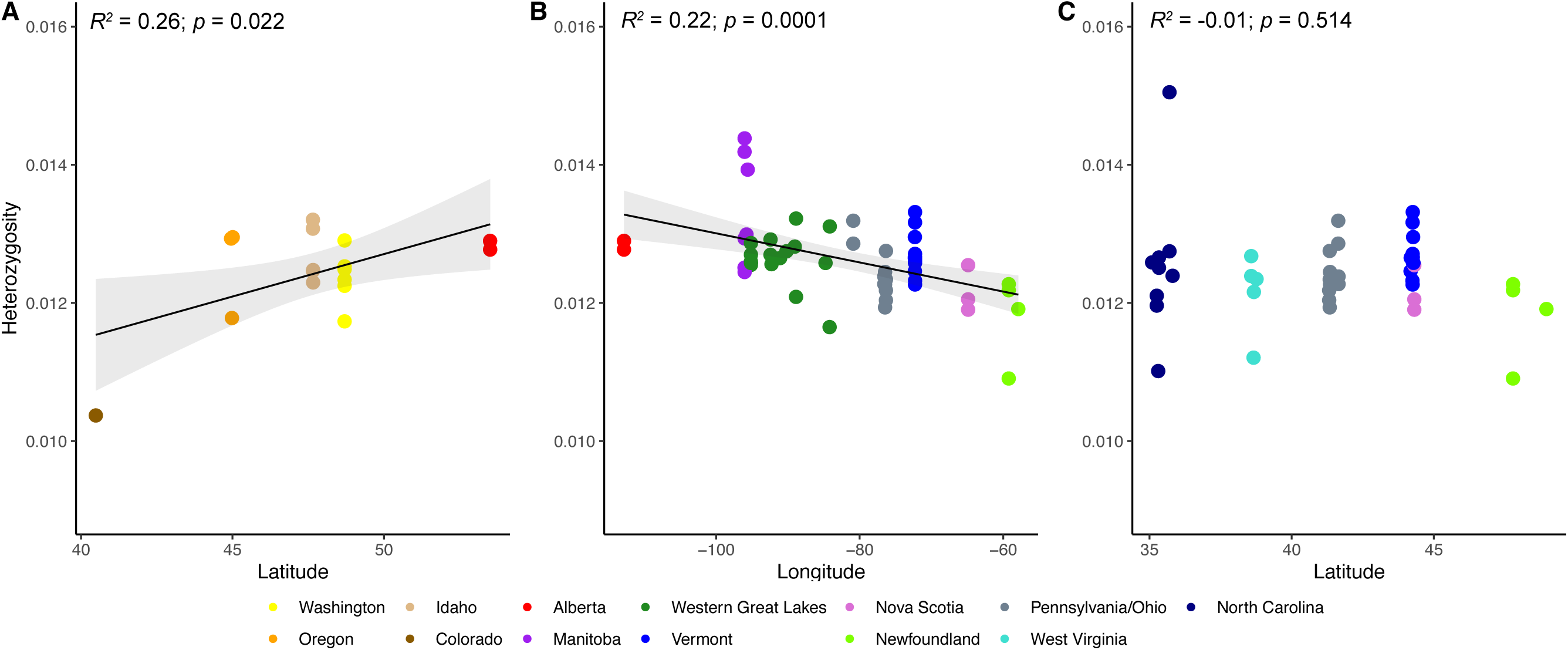
Gradients of heterozygosity across (A) the western montane region by latitude, (B) the boreal forest belt by longitude, and (C) the eastern montane region (southern Appalachians northeast to Newfoundland) by latitude. Heterozygosity scales positively with latitude across the western montane region. Heterozygosity is also significantly higher in the west than the east across the boreal forest belt. There is no significant relationship between heterozygosity and latitude across the eastern montane region. Points are colored according to their sampling region.

**Table 1.**
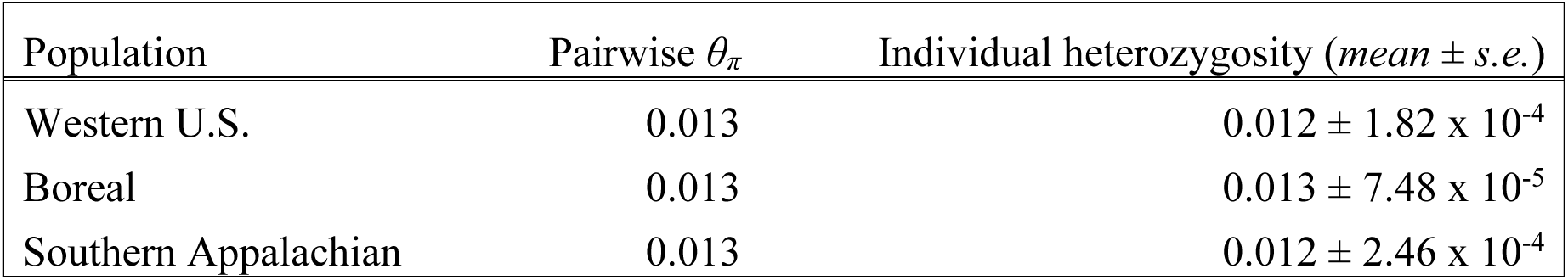
Nucleotide diversity (pairwise *θ_π_*) estimated per chromosome and individual-level heterozygosity estimated as the number of polymorphic sites divided by the total sites in each individual’s 1D site frequency spectrum.

| Population | Pairwise $\theta_\pi$ | Individual heterozygosity (*mean* ± *s.e.*) |
| --- | --- | --- |
| Western U.S. | 0.013 | $0.012 \pm 1.82 \times 10^{-4}$ |
| Boreal | 0.013 | $0.013 \pm 7.48 \times 10^{-5}$ |
| Southern Appalachian | 0.013 | $0.012 \pm 2.46 \times 10^{-4}$ |

### (c) Breeding population assignment for nonbreeding samples

We leveraged our thorough sampling of the breeding range to assess the likely breeding populations for the 4 nonbreeding samples from South America based on their location in a PCA of all individuals. One nonbreeding bird (collected in Bolivia in November) clustered with the Appalachian breeding samples from West Virginia, and the remaining three individuals (collected in Bolivia in November or Paraguay in October) clustered with boreal breeding samples (Fig. 2A). A PCA containing only the boreal individuals (Fig. 2C) suggested that one nonbreeding sample likely originated from either Manitoba or the Western Great Lakes, whereas the other two samples associate with the Western Great Lakes or the northern Appalachians (i.e., VT, PA, or OH). However, without distinct clusters in the boreal-only analysis, we refrain from confidently assigning these nonbreeding samples to breeding populations more specific than the broader boreal population.

## Discussion

We found evidence of isolation by distance across the breeding range of the Veery, a long-distance migratory songbird, as well as population clustering of the western, boreal, and southern Appalachian sampling regions. PCA revealed geographically nested patterns of genetic clustering (Fig. 2) and a pattern of genetic covariance between individual samples that decayed with geographic distance (Fig. 3). Our results suggest that, despite long-distance migratory birds’ high dispersal ability and long migratory journeys, site fidelity, which acts to lower natal and breeding dispersal distances, appears to be sufficiently strong to yield fine-scale geographic genetic structure in the absence of extrinsic barriers to dispersal.

Previous phenotypic assessments of the Veery described five subspecies based on plumage color variation associated with the following regions: (1) Newfoundland and central Quebec (*C. f. fuliginosus*), (2) the eastern United States and Canada (including all Appalachian populations; *C. f. fuscescens*), (3) the Great Plains of Canada and western Great Lakes regions (*C. f. levyi*), (4) British Columbia and the Rocky Mountains (*C. f. salicicolus*), and the (5) western United States east of the Cascade Mountains (*C.f. subpallidus*) (Heckscher et al. 2020). Our genetic results are inconsistent with the boundaries of these phenotypically described subspecies, as we found only three differentiated populations across the range of the Veery, with the southern Appalachian population the most distinct. The boreal and northern Appalachian PCA revealed that 3 of our 4 samples from Newfoundland clustered together separately from the other boreal samples, suggesting a subtle genetic difference in that sampling region. Clustering of Newfoundland samples, however, could not be quantitatively identified as a discrete population. We conclude that the genetic structure detected does not align with the phenotypically described subspecies, such that phenotypic differences are unlikely driven by historical population isolation and differentiation (Zamudio et al. 2016). Instead, subtle plumage differences across the range could reflect local selection on a small number of plumage genes without genome-wide divergence (e.g., McCormack et al. 2012) or phenotypic plasticity in response to environmental conditions (e.g., Mason and Taylor 2015; López-Rull et al. 2023).

Our data also allowed us to determine the general breeding origins of the very few wintering site genetic samples available. Understanding migratory connectivity or the geographic links between wintering, stopover, and breeding sites is critical (Webster and Marra 2005; Marra et al. 2006; Somveille et al. 2021), as conditions on the wintering grounds can have carry-over effects on breeding season fitness (Norris and Taylor 2006; Harrison et al. 2011; Ambrosini et al. 2019). Individual tracking can reveal movement patterns across the annual cycle (Stutchbury et al. 2009; Fraser et al. 2012; Batbayar et al. 2021; Rushing et al. 2021), but is both time intensive and accompanied by several challenges associated with sample size and data recovery (Ruegg et al. 2017). The Veery’s complex movements between two wintering regions in the tropics (Heckscher et al. 2011; Heckscher et al. 2015; Hobson and Kardynal 2015) add another challenge to using tracking information to identify the breeding population of an individual. Genetic data from whole-genome sequencing has been used previously to identify an individual’s population of origin (e.g., Manel et al. 2002; Nielsen et al. 2009; Hess et al. 2011; Ruegg et al. 2014) and might be a robust alternative method to tracking methods, as it is cost effective at a large scale and can be used to detect subtle breakpoints in continuous population structure (Turbek et al. 2023). We used PCA to identify putative population of origin for the four nonbreeding birds in our dataset that were likely sampled at their first wintering site. We identified one individual to be from the southern Appalachian population, whereas the other three nonbreeding birds seem to have originated from the boreal population (Fig. 2). Although the lack of distinct genetic clusters in the boreal region lowers confidence in finer-scale assignment of breeding population origins within this region, one winter sample clustered closely with several individuals sampled in the Western Great Lakes, whereas the other two samples were associated with individuals from the Northeast U.S. Most population assignment techniques use a panel of genetic markers or loci that consistently differ between distinct populations (Veale et al. 2012; Chen et al. 2018a; Sylvester et al. 2018); however, these techniques are ineffectual across wide ranges without pronounced population structure, such as the boreal forest belt for the Veery. By combining lcWGS with range-wide sampling, PCAs can detect fine-scale structure, such that regional breeding area assignment might be possible in regions with high gene flow.

We also evaluated summary statistics associated with differences in genetic diversity between populations to test whether contemporary genetic patterns reflected historic isolation of populations in glacial refugia. We expected that if the southern Rockies and southern Appalachia were glacial refugia for the Veery, then we would detect lower genetic diversity in the boreal population in comparison (Hewitt 1999; Provan and Bennett 2008). We found that the western, boreal, and Appalachian populations did not differ in any measures of genetic diversity (Table 1, Fig. 4). These results do not support predictions of southern glacial refugia (Hewitt 2004; Provan and Bennett 2008; Ralston et al. 2021).

Population-level differences in genetic diversity can also show signatures of range expansion dynamics (Provan and Bennett 2008; Peter and Slatkin 2015; Adams et al. 2023). Due to founder effects and greater geographic isolation, the expansion front is expected to have lower levels of genetic diversity compared to the core of the range (García-Ramos and Kirkpatrick 1997; Excoffier 2004; Eckert et al. 2008; Excoffier et al. 2009). By contrast, the core of the range may harbor higher genetic diversity due to prolonged population stability (Provan and Bennett 2008; van Els et al. 2012). We found that individual heterozygosity was positively related to latitude across the western montane region and negatively with longitude across the boreal forest belt (Fig. 5). These patterns do not align with expected gradients of genetic diversity if contemporary genetic patterns reflect northward postglacial expansion as the glaciers retreated (Miller et al. 2020; Adams et al. 2023). However, comparable or higher genetic diversity has also been observed at the leading expansion front (Vandepitte et al. 2017; Wang et al. 2017; Bors et al. 2019) likely due to continued high gene flow with the core population (Miller et al. 2020; Adams et al. 2023). Expansions that occur at a rapid pace are also likely to retain higher heterozygosity at the expansion front (Goodsman et al. 2014). Therefore, our heterozygosity results may alternatively provide weak support for rapid expansions out of western and northeastern refugia. The directionality index, however, was close to zero between all pairwise comparison (*ψ* < 0.03), suggesting that the data might better fit an isolation by distance rather than expansion model (Peter and Slatkin 2013; Adams et al. 2023). Ultimately, our results do not provide compelling evidence for a glacial refugium in Newfoundland or the southern Rockies, because the subtle patterns found are also consistent with continuous processes of gene flow between populations across the range.

In conclusion, we were able to resolve geographic fine-scale genetic structure in the Veery despite the high dispersal potential in this species, and we observed evidence for both continuous and discontinuous structure across the range. Given the fine-scale resolution achieved through low-coverage, whole-genome sequencing and dense sampling across the range, we were also able to assign regions of origin to individuals collected on their wintering grounds, which has important implications for assessing migratory connectivity at a larger scale than enabled by traditional tracking methods. Finally, based on the patterns of population differentiation and genetic diversity in this species, we conclude that gene flow, isolation by distance, and site fidelity likely play a more important role in shaping current population genetic structure and diversity in this species than historic isolation.

## Acknowledgments

For providing tissue and blood samples, we thank the American Museum of Natural History (Brian Smith, Joel Cracraft, Paul Sweet, Peter Capainolo, Tom Trombone), New York State Museum (Jeremy Kirchman), University of Washington Burke Museum (Sharon Birks), Royal Alberta Museum (Jocelyn Hudon), University of Kansas Biodiversity Institute (Town Peterson, Robert Moyle, Mark Robbins), Cleveland Museum of Natural History (Courtney Brennan), University of Michigan Museum of Zoology (Brett Benz) and the University of Alaska Museum. For assistance in the field, we thank Courtney Brennan, Brett Benz, Susanna Campbell, Shane DuBay, Laura Gooch, Eric Gulson-Castillo, Ethan Gyllenhaal, Heather Skeen, Vera Ting, and Brian Weeks. We also thank Kristen Ruegg and Teia Schweizer for training and support in their whole-genome library preparation methods. We thank Nicole Adams, Gideon Bradburd, Zach Hancock, and Leonard Jones for discussion and feedback. Next-generation sequencing for this project was carried out in the Advanced Genomics Core at the University of Michigan. This research was also supported in part through computational resources and services provided by Advanced Research Computing (ARC), a division of Information and Technology Services (ITS) at the University of Michigan, Ann Arbor.

## Ethics statement

For permits to collect specimens in the field, we thank the United States Fish and Wildlife Service, United States Forest Service, Michigan Department of Natural Resources, Minnesota Department of Natural Resources, Colorado Department of Natural Resources, Idaho Department of Fish and Game, North Carolina Wildlife Resources Commission, Ohio Division of Wildlife, Oregon Department of Fish and Wildlife, Pennsylvania Game Commission, Vermont Fish and Wildlife Department, Vermont Agency of Natural Resources, West Virginia Department of Natural Resources, Canadian Wildlife Service of Environment and Climate Change Canada, and Manitoba Fish and Wildlife. Field sampling was approved by the University of Michigan Animal Care and Use Committee (# PRO00010608).

## Author Contributions

- AWJ, AAK, TMP, and BMW conceived the idea of the study.
- AAK, TMP, and BMW developed methodology.
- AAK and BMW analyzed the data.
- AAK and BMW wrote the original draft of the manuscript.
- TMP, AWJ and KW reviewed and edited the manuscript.
- AWJ, KW, and BMW contributed substantial materials or funding for the study.

