## Supplementary Materials for "How Veeries vary: Whole genome sequencing resolves fine-scale genetic structure in a long-distance migratory bird, *Catharus fuscescens*"

**Table S1.** Summary table of samples used in analyses, which includes institution, species, catalogue number, sex, collection date, locality, and latitude/longitude data for each sample. Acronyms for institution names are as follows: AMNH=American Museum of Natural History; CMNH= Cleveland Museum of Natural History; KU = University of Kansas Biodiversity Institute; NYSM= New York State Museum; RAM= Royal Alberta Museum; UAM= University of Alaska Museum; UMMZ= University of Michigan Museum of Zoology; UWBM= University of Washington Burke Museum.

| Institution | Catalog Number | Sex | Date | State/ Province | County/ District | Latitude | Longitude |
|---|---|---|---|---|---|---|---|
| **AMNH** | DOT-6090 | U | 11/8/00 | Bolivia | Cordillera | -19.77 | -62.7 |
| **AMNH** | DOT-6123 | U | 11/12/00 | Bolivia | Cordillera | -19.77 | -62.7 |
| **AMNH** | DOT-6134 | M | 11/13/00 | Bolivia | Cordillera | -19.77 | -62.7 |
| **CMNH** | 72522 | M | 6/5/12 | Colorado | Routt | 40.487682 | -107.15524 |
| **CMNH** | 72523 | M | 6/5/12 | Colorado | Routt | 40.487682 | -107.15524 |
| **CMNH** | 72576 | M | 6/8/12 | Colorado | Routt | 40.487682 | -107.15524 |
| **CMNH** | 73127 | M | 6/16/14 | Idaho | Shoshone | 47.65335 | -116.07384 |
| **CMNH** | 73129 | M | 6/16/14 | Idaho | Shoshone | 47.65335 | -116.07384 |
| **CMNH** | 73145 | M | 6/16/14 | Idaho | Shoshone | 47.66053 | -116.05981 |
| **CMNH** | 73146 | M | 6/16/14 | Idaho | Shoshone | 47.66053 | -116.05981 |
| **CMNH** | 73147 | M | 6/16/14 | Idaho | Shoshone | 47.65335 | -116.07384 |
| **CMNH** | 72623 | M | 6/25/11 | Michigan | Oscoda | 44.79176 | -84.17626 |
| **CMNH** | 73720 | M | 6/25/11 | Michigan | Oscoda | 44.7654 | -84.19229 |
| **CMNH** | 76949 | M | 6/7/18 | Michigan | Gogebic | 46.16035 | -89.04452 |
| **CMNH** | 76972 | M | 6/10/18 | Michigan | Houghton | 46.60996 | -88.86445 |
| **CMNH** | 71983 | M | 6/10/11 | Minnesota | Hubbard | 47.28239 | -95.15358 |
| **CMNH** | 71984 | M | 6/10/11 | Minnesota | Hubbard | 47.28239 | -95.15358 |
| **CMNH** | 72265 | M | 6/13/11 | Minnesota | Hubbard | 47.30525 | -95.14462 |
| **CMNH** | 72269 | M | 6/16/11 | Minnesota | St Louis | 48.0046 | -92.42556 |
| **CMNH** | 72270 | M | 6/16/11 | Minnesota | St Louis | 48.0046 | -92.42556 |
| **CMNH** | 72271 | M | 6/11/11 | Minnesota | Hubbard | 47.25461 | -95.15334 |
| **CMNH** | 72272 | M | 6/11/11 | Minnesota | Hubbard | 47.25461 | -95.15334 |
| **CMNH** | 72280 | M | 6/15/11 | Minnesota | St Louis | 48.11102 | -92.28787 |

| | | | | | | | |
|---|---|---|---|---|---|---|---|
| **CMNH** | 75061 | M | 6/11/17 | Minnesota | Cook | 47.972 | -90.205 |
| **CMNH** | 75134 | M | 6/15/17 | Minnesota | Lake | 47.828 | -91.086 |
| **CMNH** | 70513 | M | 5/21/07 | North Carolina | Buncombe | 35.70034479 | -82.379634 |
| **CMNH** | 70514 | M | 5/21/07 | North Carolina | Buncombe | 35.70034479 | -82.379634 |
| **CMNH** | 70515 | M | 5/21/07 | North Carolina | Buncombe | 35.70034479 | -82.379634 |
| **CMNH** | 70573 | M | 5/25/07 | North Carolina | Buncombe | 35.08973 | -82.34804 |
| **CMNH** | 70642 | M | 5/25/07 | North Carolina | Buncombe | 35.80973 | -82.34804 |
| **CMNH** | 70894 | M | 5/31/08 | North Carolina | Transylvania | 35.25308 | -82.91874 |
| **CMNH** | 70895 | M | 5/31/08 | North Carolina | Transylvania | 35.25308 | -82.91874 |
| **CMNH** | 70896 | M | 5/31/08 | North Carolina | Transylvania | 35.25308 | -82.91874 |
| **CMNH** | 70897 | M | 5/30/08 | North Carolina | Haywood | 35.32986 | -82.90807 |
| **CMNH** | 70898 | M | 5/30/08 | North Carolina | Haywood | 35.32986 | -82.90807 |
| **CMNH** | 70899 | M | 5/31/08 | North Carolina | Haywood | 35.376592 | -82.789371 |
| **CMNH** | 70900 | M | 5/29/08 | North Carolina | Buncombe | 35.81214 | -82.356 |
| **CMNH** | 70901 | M | 5/31/08 | North Carolina | Transylvania | 35.25308 | -82.91874 |
| **CMNH** | 71885 | M | 5/30/08 | North Carolina | Haywood | 35.30361 | -82.90871 |
| **CMNH** | 70351 | M | 6/12/06 | Ohio | Ashtabula | 41.63547871 | -80.892671 |
| **CMNH** | 70352 | M | 6/9/06 | Ohio | Ashtabula | 41.63547871 | -80.892671 |
| **CMNH** | 70374 | M | 6/14/06 | Ohio | Ashtabula | 41.63547871 | -80.892671 |
| **CMNH** | 70638 | M | 6/15/07 | Ohio | Ashtabula | 41.63547871 | -80.892671 |
| **CMNH** | 70666 | M | 6/14/07 | Ohio | Ashtabula | 41.63547871 | -80.892671 |
| **CMNH** | 73047 | M | 6/7/14 | Oregon | Baker | 44.98092 | -117.37164 |
| **CMNH** | 73073 | M | 6/9/14 | Oregon | Baker | 44.95285 | -117.33763 |
| **CMNH** | 73075 | M | 6/8/14 | Oregon | Union | 45.00677 | -117.57866 |
| **CMNH** | 73077 | M | 6/9/14 | Oregon | Union | 45.00677 | -117.57866 |
| **CMNH** | 71021 | M | 6/13/08 | Pennsylvania | Sullivan | 41.34671 | -76.46123 |
| **CMNH** | 71025 | M | 6/12/08 | Pennsylvania | Sullivan | 41.33601 | -76.462 |
| **CMNH** | 71026 | M | 6/12/08 | Pennsylvania | Sullivan | 41.34671 | -76.46123 |
| **CMNH** | 71063 | M | 6/12/08 | Pennsylvania | Sullivan | 41.34671 | -76.46123 |

| | | | | | | | |
|---|---|---|---|---|---|---|---|
| **CMNH** | 71064 | M | 6/11/08 | Pennsylvania | Bradford | 41.64682 | -76.57269 |
| **CMNH** | 71070 | M | 6/11/08 | Pennsylvania | Bradford | 41.64682 | -76.57269 |
| **CMNH** | 71071 | M | 6/11/08 | Pennsylvania | Bradford | 41.64682 | -76.57269 |
| **CMNH** | 71188 | M | 6/14/08 | Pennsylvania | Sullivan | 41.33508 | -76.33997 |
| **CMNH** | 71189 | M | 6/14/08 | Pennsylvania | Sullivan | 41.33508 | -76.33997 |
| **CMNH** | 71221 | M | 6/14/08 | Pennsylvania | Sullivan | 41.33508 | -76.33997 |
| **CMNH** | 71331 | M | 6/13/08 | Pennsylvania | Sullivan | 41.31794 | -76.34454 |
| **CMNH** | 71420 | M | 6/13/08 | Pennsylvania | Sullivan | 41.39627 | -76.42828 |
| **CMNH** | 71930 | M | 6/13/08 | Pennsylvania | Sullivan | 41.31794 | -76.34454 |
| **CMNH** | 71777 | M | 6/21/08 | Vermont | Caledonia | 44.25215 | -72.28686 |
| **CMNH** | 71778 | M | 6/19/08 | Vermont | Caledonia | 44.2571 | -72.28785 |
| **CMNH** | 71779 | M | 6/19/08 | Vermont | Caledonia | 44.2571 | -72.28785 |
| **CMNH** | 71780 | M | 6/19/08 | Vermont | Caledonia | 44.2571 | -72.28785 |
| **CMNH** | 71836 | M | 6/21/08 | Vermont | Caledonia | 44.25215 | -72.28686 |
| **CMNH** | 71993 | M | 6/21/08 | Vermont | Caledonia | 44.25215 | -72.28686 |
| **CMNH** | 71994 | M | 6/21/08 | Vermont | Caledonia | 44.25215 | -72.28686 |
| **CMNH** | 71996 | M | 6/18/08 | Vermont | Caledonia | 44.27053 | -72.30969 |
| **CMNH** | 72004 | M | 6/20/08 | Vermont | Orange | 44.18084 | -72.30788 |
| **CMNH** | 72147 | M | 6/21/08 | Vermont | Caledonia | 44.28233 | -72.28834 |
| **CMNH** | 72334 | M | 6/20/08 | Vermont | Orange | 44.186 | -72.32238 |
| **CMNH** | 72652 | M | 6/21/08 | Vermont | Caledonia | 44.25215 | -72.286886 |
| **CMNH** | 72663 | M | 6/21/08 | Vermont | Caledonia | 44.25215 | -72.28686 |
| **CMNH** | 72664 | M | 6/19/08 | Vermont | Caledonia | 44.24557 | -72.28219 |
| **CMNH** | 72675 | M | 6/21/08 | Vermont | Caledonia | 44.24557 | -72.28219 |
| **CMNH** | 73048 | M | 6/7/14 | Oregon | Baker | 44.98198 | -117.37054 |
| **CMNH** | 70903 | M | 6/5/08 | West Virginia | Randolph | 38.66839 | -79.90349 |
| **CMNH** | 70904 | M | 6/5/08 | West Virginia | Randolph | 38.66839 | -79.90349 |
| **CMNH** | 70905 | M | 6/5/08 | West Virginia | Randolph | 38.66839 | -79.90349 |
| **CMNH** | 70906 | M | 6/5/08 | West Virginia | Randolph | 38.69173 | -79.88634 |

| | | | | | | | |
|---|---|---|---|---|---|---|---|
| **CMNH** | 70907 | M | 6/5/08 | West Virginia | Randolph | 38.69173 | -79.88634 |
| **CMNH** | 70908 | M | 6/5/08 | West Virginia | Randolph | 38.69173 | -79.88634 |
| **CMNH** | 70909 | M | 6/5/08 | West Virginia | Randolph | 38.69173 | -79.88634 |
| **CMNH** | 71056 | M | 6/6/08 | West Virginia | Pocahontas | 38.57568 | -79.82653 |
| **CMNH** | 71059 | M | 6/6/08 | West Virginia | Pocahontas | 38.57568 | -79.82653 |
| **CMNH** | 71061 | M | 6/5/08 | West Virginia | Randolph | 38.65794 | -79.90945 |
| **CMNH** | 71065 | M | 6/5/08 | West Virginia | Randolph | 38.65794 | -79.90945 |
| **CMNH** | 71066 | M | 6/5/08 | West Virginia | Randolph | 38.65794 | -79.90945 |
| **CMNH** | 71068 | M | 6/3/08 | West Virginia | Randolph | 38.76311 | -79.8482 |
| **CMNH** | 71069 | M | 6/2/08 | West Virginia | Randolph | 38.65065 | -79.91332 |
| **CMNH** | 71072 | M | 6/6/08 | West Virginia | Pocahontas | 38.64029 | -79.75553 |
| **KU** | 88484 | M | 10/30/96 | Paraguay | Concepción | -22.6666667 | -57.35 |
| **NYSM** | zt-1282 | F | 7/11/15 | Newfoundland | C.D. No. 4 | 47.7831 | -59.23317 |
| **NYSM** | zt-1291 | M | 7/11/15 | Newfoundland | C.D. No. 4 | 47.7831 | -59.23317 |
| **NYSM** | zt-1292 | M | 7/11/15 | Newfoundland | C.D. No. 4 | 47.7831 | -59.23317 |
| **RAM** | Z02.10.1 | M | 5/30/02 | Alberta | C.D. No. 10 | 53.504167 | -112.83944 |
| **RAM** | Z02.16.7 | M | 7/5/02 | Alberta | C.D. No. 10 | 53.506111 | -112.8425 |
| **UAM** | 19845 | M | 6/17/93 | Newfoundland | C.D. No. 5 | 48.95941987 | -57.897909 |
| **UAM** | 13414 | M | 6/26/93 | Nova Scotia | Queens | 44.3116926 | -64.889 |
| **UAM** | 13416 | M | 6/26/93 | Nova Scotia | Queens | 44.3116926 | -64.889 |
| **UAM** | 19846 | M | 6/25/93 | Nova Scotia | Queens | 44.3116926 | -64.889 |
| **UAM** | 19847 | M | 6/25/93 | Nova Scotia | Queens | 44.3116926 | -64.889 |
| **UMMZ** | 247374 | M | 6/9/19 | Manitoba | C.D. No. 19 | 50.41793 | -95.60412 |
| **UMMZ** | 247375 | M | 6/5/19 | Manitoba | C.D. No. 1 | 49.549852 | -95.770257 |
| **UMMZ** | 247376 | M | 6/6/19 | Manitoba | C.D. No. 1 | 49.769601 | -96.043321 |
| **UMMZ** | 247377 | M | 6/6/19 | Manitoba | C.D. No. 1 | 49.769601 | -96.043321 |
| **UMMZ** | 247378 | M | 6/6/19 | Manitoba | C.D. No. 1 | 49.791 | -96.034916 |
| **UMMZ** | 247379 | M | 6/6/19 | Manitoba | C.D. No. 1 | 49.791 | -96.034916 |
| **UMMZ** | 247380 | M | 6/6/19 | Manitoba | C.D. No. 1 | 49.791 | -96.034916 |

| | | | | | | | |
|---|---|---|---|---|---|---|---|
| **UMMZ** | 245925 | F | 6/12/18 | Michigan | Houghton | 46.660097 | -88.86901 |
| **UMMZ** | 248008 | M | 6/8/21 | Michigan | Emmet | 45.59466 | -84.75956 |
| **UMMZ** | 248009 | M | 6/7/21 | Michigan | Emmet | 45.569819 | -84.967727 |
| **UMMZ** | 248010 | M | 6/11/21 | Michigan | Presque Isle | 45.475582 | -84.169084 |
| **UMMZ** | 248011 | F | 6/10/21 | Michigan | Presque Isle | 45.475582 | -84.169084 |
| **UWBM** | 62067 | F | 6/2/99 | Washington | Okanogan | 48.7 | -119.7 |
| **UWBM** | 62068 | F | 6/2/99 | Washington | Okanogan | 48.7 | -119.7 |
| **UWBM** | 62071 | F | 6/4/99 | Washington | Okanogan | 48.7 | -119.7 |
| **UWBM** | 62078 | M | 6/3/99 | Washington | Okanogan | 48.7 | -119.7 |
| **UWBM** | 62084 | F | 6/2/99 | Washington | Okanogan | 48.7 | -119.7 |
| **UWBM** | 62136 | M | 6/2/99 | Washington | Okanogan | 48.7 | -119.7 |
| **UWBM** | 62144 | M | 6/2/99 | Washington | Okanogan | 48.7 | -119.7 |
| **UWBM** | 62151 | M | 6/2/99 | Washington | Okanogan | 48.7 | -119.7 |
| **UWBM** | 79282 | F | 6/4/99 | Washington | Okanogan | 48.7 | -119.7 |

**Table S2.** Parameters used in ANGSD to estimate metrics. Parameter definitions were summarized from the ANGSD website.

| Parameter | Definition | Genotype likelihoods | Site allele frequency | Individual-level heterozygosity |
|---|---|---|---|---|
| **uniqueOnly** | Remove reads that have multiple best hits | 1 | 1 | 1 |
| **remove_bads** | Removes read with a flag above 255 | 1 | 1 | 1 |
| **only_proper_pairs** | Include pairs of read with both mates mapped correctly | 1 | 1 | 1 |
| **minMapQ** | Minimum mapQ quality | 30 | 30 | 30 |
| **minQ** | Minimum base quality score | 30 | 30 | 30 |
| **SNP_pval** | Only use sites with a p-value less than [value] | 0.05 | | |
| **minMaf** | Only use sites with a minor allele frequency above [value] | 0.05 | | |
| **GL** | Estimate genotype likelihoods | 2 | 2 | 2 |
| **doMajorMinor** | Define what is the major and minor allele | 1 | | |
| **doMaf** | Estimate allele frequencies | 1 | | |
| **doGlf** | Define the output file format | 2 | | |
| **doCounts** | Estimate frequencies of the different bases | 1 | | |
| **doSaf** | Estimate site allele frequency likelihoods | | 1 | 1 |
| **doHWE** | Test for HWE based on genotype likelihoods | 1 | | |
| **dumpCounts** | Outputs depth of each individual | 2 | | |
| **doDepth** | Outputs distribution of sequencing depths | 1 | | |
| **dosnpstat** | Report stats without filtering sites | 1 | | |

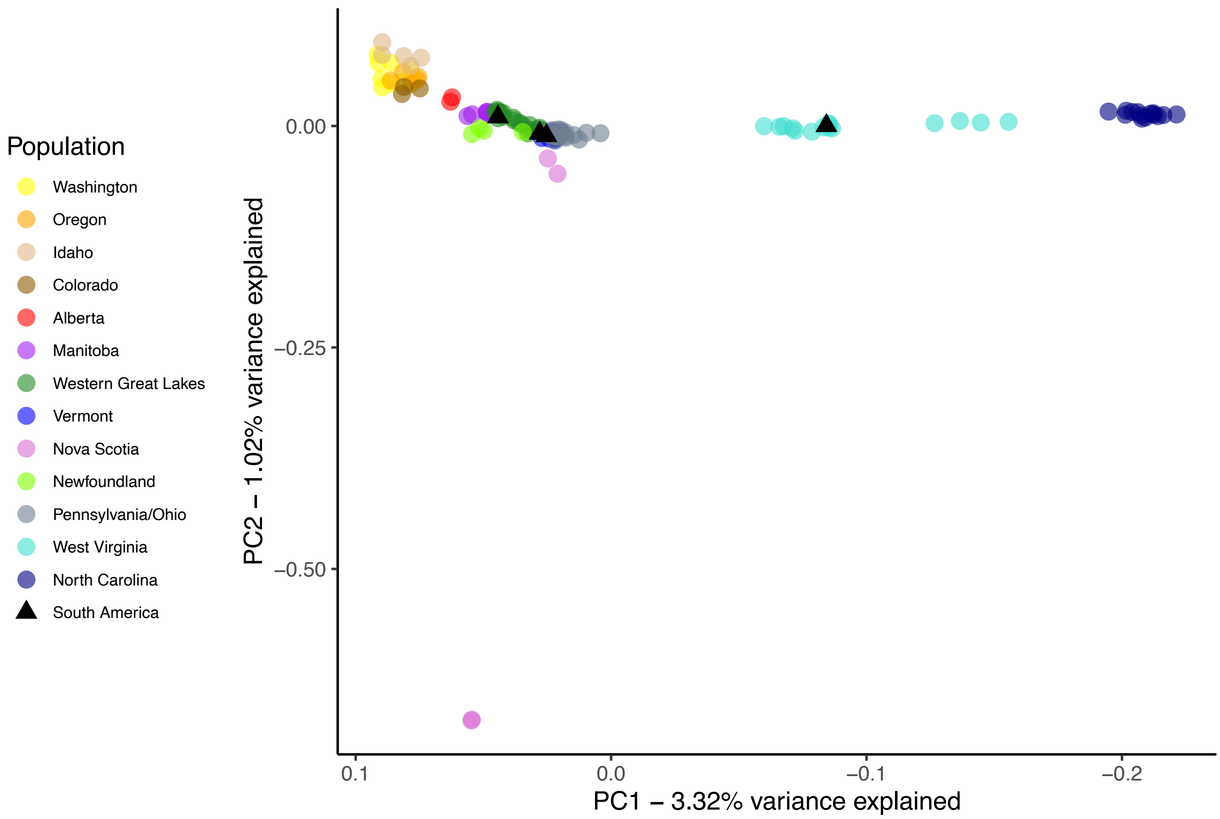


**Figure S1.** Principal Component Analysis (PCA) for all samples including both Nova Scotia samples that were visual outliers in the plot. Nonbreeding birds collected in the South America are displayed as black triangles to identify potential population of origin. The x-axis is reversed so that points are displayed west to east. The southern Appalachian (West Virginia and North Carolina) were supported as a distinct population from all other samples.
